# Genetic and histological correlation between the musk gland and skin of Chinese forest musk deer

**DOI:** 10.1101/2022.10.15.512364

**Authors:** Long Li, Heran Cao, Jinmeng Yang, Yuxuan Ma, Tianqi Jin, Yang Wang, Zhenpeng Li, Yining Chen, Huihui Gao, Chao Zhu, Tianhao Yang, Yalong Deng, Fangxia Yang, Wuzi Dong

## Abstract

Chinese forest musk deer (FMD, *Moschus berezovskii*) glands have the ability to secrete musk, which plays an important role in attracting females during the breeding season. Sebaceous glands (SGs) are exocrine skin glands associated with hair follicles that continuously release a mixture of sebum, lipids and cellular debris, by holocrine secretion. Both the musk glands and the skin tissues of the FMD contain abundant sebaceous glands, and *Sox9, Caveolin1*, and *Androgen receptor* (*AR*) are all involved in the regulation of sebum secretion by the sebaceous glands. However, there are fewer studies on the correlation between skin and musk glands and the expression of *Sox9, Caveolin1*, and *AR* in the musk glands and skin tissue of FMD. To address this gap, we analyzed biochemical data from FMD skin tissues and musk glands using transcriptomic data, hematoxylin-eosin (HE) staining, western blotting (WB), immunohistochemistry (IHC), tissue dissection, and RT-qPCR. Anatomical results show that only adult male FMD had complete glandular part and sachets, while 4-month-old FMD do not have well-developed sachets. Transcriptomic data showed that 88.24% of genes were co-expressed in the skin and musk glands tissues of FMD. The WB, IHC, and RT-qPCR results showed that the genes involved in regulating sebum secretion, *Sox9, Caveolin1*, and *AR* were expressed in the skin tissues and musk glands. In summary, skin tissues and musk glands tissue have a strong correlation, and *Sox9, Caveolin1*, and *AR* may play important roles in skin tissues and musk glands tissue.

## Introduction

Mammalian pheromones differ from animal scent signatures consisting of complex and variable mixtures, which possess function-specific, species-specific responses(Shirasu *et al*. 2020). Pheromones are synthesized and released in animals and act in the outside world to communicate with other organisms of the same species and have been extensively studied in insects, mammals, and humans (Varendi and Porter 2001; Slessor *et al*. 2005). The three primary appendages of skin tissue, namely sebaceous glands, eccrine sweat glands, hair follicles, which are derived from the developing epidermis (Saga 2002; Sotiropoulou and Blanpain 2012; Biggs and Mikkola 2014; Fuchs 2016).

The musk secretion of Chinese FMD is important for breeding, territory identification, and medicinal purposes, and the main factor regulating the secretion of adult males are the level of androgens (Jiang *et al*. 2021; Yang *et al*. 2021). The previous analysis of the chemical composition of musk found that musk mainly contains fatty acids, cholesterol, esters, alcohols, and cyclic ketones, of which fatty acids account for about 71.55%, and the remaining components are cholesterol (9.31%) and other organic substances (Li *et al*. 2017; Zhang *et al*. 2021b). Musk secreted by adult male FMD has a strong smelling, which is used to mark territory and attract females (Fan *et al*. 2018; Li *et al*. 2018). The musk gland is an exocrine gland, and only the sexually mature male FMD can secrete musk (Sokolov *et al*. 1987). The gland’s epithelial cells secrete musk in the form of an apocrine manner, which enters the sachets via a duct (Chen *et al*. 2018).

The sebum secreted by the sebaceous glands in the neck and ducts of the bursa enters the sachets, where the immature musk and sebum mature gradually for about two months (Chen *et al*. 2018; Lv *et al*. 2022). Most sebaceous glands are associated with hair follicles and form follicular sebaceous gland units that produce sebum through differentiation and disintegration of fully mature sebaceous cells (Thody and Shuster 1989; Downie *et al*. 2004; Smith and Thiboutot 2008; Zouboulis *et al*. 2008; Schneider and Paus 2010). *Sox9* is located in keratinocytes, sebaceous glands, sweat glands, and melanocytes (Vidal *et al*. 2008; Shi *et al*. 2013; Shakhova *et al*. 2015). Studies have shown that *Sox9* can promote the proliferation, differentiation and sebaceous secretion of sebaceous gland cells (Shi *et al*. 2017). *Sox9* is essential for the development of sebaceous glands, can participate in regulating hair follicle stem cell differentiation, and is of great significance for the morphogenesis of hair follicles and sebaceous glands (Nowak *et al*. 2008; Schneider and Paus 2010; TÓth *et al*. 2011). Caveolin1 is highly expressed in many cells and can interact with multiple proteins, which is the signal transduction hub of cell signaling molecules (Boscher and Nabi 2012; Fridolfsson *et al*. 2014). Studies have shown the ability of *Caveolin1* to promote cell differentiation in a specific direction in a variety of cells (Fu *et al*. 2012; Codenotti *et al*. 2016). *Caveolin1* plays an important role in regulating lipid metabolism, especially adipose tissue (Bastiani *et al*. 2009). The androgen receptor (AR) is a major member of the steroid receptor superfamily and is expressed in many skin cells such as hair follicles, sebaceous glands, and sweat glands (Barrault *et al*. 2015; Davey and Grossmann 2016). The secretion of sebum is closely related to the function of the sebaceous glands, and androgens regulate the function of the sebaceous glands through androgen receptors, of which dihydrotestosterone is the most effective androgen (Huang *et al*. 2018; Duarte *et al*. 2019). The *Sox9*/ *AR*/ Wnt/ β-catenin signaling pathway can express target genes for cell viability, proliferation, and differentiation (Khurana and Sikka 2019). AR binds to Caveolin1 depending on the plasma membrane through palmitoylation of cysteine residues, and testosterone induces AR to move toward the cell membrane site and bind to Caveolin1 (Bennett *et al*. 2009). Therefore, we hypothesize that the musk glands of FMD, as a specialized secretory gland with some similarity with skin tissues, may also be regulated by *Sox9, Caveolin1*, and *AR* in the development and secretion process. However, this is a gap about the correlation between skin tissue and musk glands, as well as the expression of regulatory genes in sebaceous glands and hair follicles. To clarify the correlation between skin tissue and musk gland in development period, we anatomy male FMD at different periods (4-months-old and adulthood), and selected the developing 4-month-old FMD as the research object. RNA-seq, immunohistochemistry (IHC), Western blot (WB), and RT-qPCR were used to study the musk glands, abdomen skin, and back skin of 4-month-old FMD.

## Materials and methods

### Animal and ethical

All experimental animal procedures were reviewed and approved by the relevant institutional Ethical Committee of Northwest A&F University (EAMC/2020−12).

### Anatomical comparison

To find suitable subjects for this study, we selected Chinese forest musk deer during before sexual maturity, and after sexual maturity as the subjects. The FMDs from these two time periods were dissected to compare the spatial structure of the musk gland and skin tissue. In particular, the FMDs in this study consisted of one 4-month-old musk deer and one adult male musk deceased accidentally, all collected from breeding farms with breeding privileges.

### Tissue sample collection and cryopreservation

According to the guidance of the ethical committee, experimental animals were collected. Immediately after FMD died, the skin on the back (n=3), the abdominal skin around the penile mouth (n=3) and musk glands (n=3) were collected. Additionally, tissues of the musk gland and skin were collected for Immunohistochemistry (IHC), western blot, and transcriptome sequencing. Samples were trimmed using scissors into approximately 1-cm^3^ pieces and subsequently fixed with 4% paraformaldehyde for HE and IHC. Using PBS-washed tissues to remove blood, immediately transfer the samples into new RNase-free tubes and store them at -80°C before extracting total tissue RNA and protein.

### Total RNA extraction and synthesis cDNA

The total RNA of each tissue was extracted by the TRIzol reagent (Invitrogen, CA, USA). Nanodrop 2000 spectrophotometer (ThermoFisher Scientific, DE, USA) device for RNA concentration and purity testing (OD 260/280 ratio > 1.8), and the next 1μg of total eligible RNA was reverse transcribed using the cDNA synthesis kit (Code No. 6110A, Takara, Japan) for quantitative real-time PCR (RT-qPCR). All experimental steps were performed according to the manufacturer’s operating instructions.

### Total RNA extraction and RNA-sequencing

Total RNA was extracted from abdominal skin, back skin, and musk glands tissue using Trizol reagent. The purity and content of total RNA were quantified using Bioanalyzer 2100 Spectrophotometers and RNA 1000 Nano LabChip Kit (Agilent, CA, USA). Total RNA was used for subsequent experiments if the total RNA samples met the following criteria: RNA 28S: 18S ≥ 1.5: and RNA integrity number (RIN) ≥ 7.0. RNA sequencing and sequencing libraries were performed by Lianchuan Biotech (Hangzhou, China).

### RNA-sequencing bioinformatic analysis

Low-quality sequencing data were filtered out by SOAPnuke (v1.5.2) (Li *et al*. 2008). To evaluate the sequence quality, according to the clean data’s Q20, Q30, and GC content (Andrews 2014). We used the Trinity (version 2.4.0) (Haas *et al*. 2013) tool for transcriptome de novo assembly. After assembly, TransDecoder (version 3.0.1; http://transdecoder.sourceforge.net/) was used to predict the coding domain sequences (CDSs). To obtain high-quality non-redundant CDSs, those encoding sequences less than 100 amino acids were removed, and the assembled unigenes data were annotated against the non-redundant (NR) protein database (https://ftp.cbi.nlm.nih.gov/blast/db), the Gene Ontology (GO) database for functional enrichment analysis (http://geneontology.org), the Pfam database (https://pfam.xfam.org), Genomes (KEGG) database for signaling pathway analysis (https://www.genome.jp/kegg), the SwissProt database (https://www.uniprot.org/), the eggNOG database (http://eggnogdb.embl.de) and the Kyoto Encyclopedia of Genes, utilizing DIAMOND (version 0.7.12) with a threshold of E-value < 10^−5^ (Buchfink *et al*. 2015).

### Gene quantification by RT-qPCR

The mRNA expression levels of candidate genes were detected by RT-qPCR using SYBR Green I chemical regent. The RT-qPCR was carried out in a 96-well plate, and *GAPDH* was used as an endogenous reference gene. Using the specific primers to amplify the candidate genes, gene primers designed by Primer Premier 6.0 (Premier Biosoft, CA, USA) and synthesized by Tsingke Biotech (Beijing, China), and gene primer sequences are shown in Table 1. The RT-qPCR amplification conditions were comprised of an initial denaturation step of 5 min at 95 °C, followed by 40 cycles of denaturation step, (i) 95 °C for 10 sec, (ii) annealing step, 60 °C for 30 sec; and (iii) extension step, 72°C for 10 sec. The melt curve requires an additional heating step at 95 °C for 15 sec, 60 °C for 60 sec, and 95 °C for 15 sec. Comparative threshold cycles (CTs) are calculated for each target gene expression level. The *GAPDH* expression level was used to normalize the target gene and calculate target gene expression using the mathematical principle: 2^−ΔΔCt^.

**Table 1.**
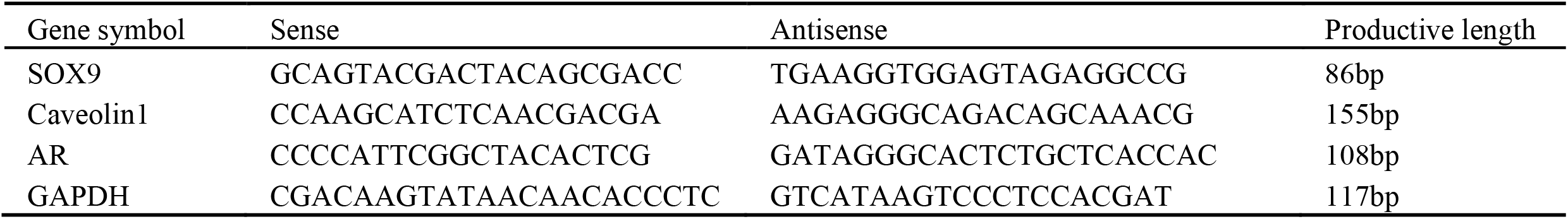
the primers information of sox9, AR, Caveolin1 for RT-qPCR

### Hematoxylin-Eosin staining and Immunohistochemistry

Collected back skin, abdominal skin, musk glandular tissue, and all obtained tissue samples were embedded in paraffin after being routinely fixed in 4% paraformaldehyde. All paraffin blocks were cut by transection cutting, and 5-µm series sections were mounted on slides. A portion of these tissue sections was used for Hematoxylin-Eosin (HE) staining, and the rest were used for immunohistochemical (IHC) experiments. In brief, the protocol of HE staining mainly includes deparaffinization, rehydration, hematoxylin, and eosin staining. Tissue morphology was observed under the microscope after tissue sealing. The IHC method was used to identify the locations of Sox9, Caveolin1, and AR in the back skin, abdominal skin, and musk glandular tissue. Dewaxing, rehydration, permeation, and antigen repair were the main components of the IHC experimental procedure. Next, use 3 % hydrogen peroxide (H_2_O_2_) incubated for 10 min to block the endogenous hydroperoxides activity. After being blocked with 5% bovine serum albumin (BSA) for 2 hours, all tissue sections were washed three times for 5 minutes each. The primary antibodies for Sox9 (1:200) (Cat. No. ab185966; Abcam; Host species: Rabbit), Caveolin1 (1:100) (Cat. No. sc-53564; Santa; Host species: Mouse), AR (1:100) (Cat. No. sc-7305; Santa; Host species: Mouse) were diluted by Dako Antibody Diluent. Then tissue sample sections were incubated with primary antibody overnight in a 4° refrigerator. At the same time, secondary antibody labeling was incubated for 1 hour at 25°C according to the protocol of the SPlink Detection Kit (Zsbio Commerce Store, Beijing, China). Finally, after DAB staining, tissue sections in the darkness for 5 min and counterstained with hematoxylin reagent for 3 min.

### Western blot analysis

Total tissue protein was extracted from the back skin, abdominal skin, musk gland, respectively. The tissues were homogenized in RIPA (RIPA: PMSF = 100:1, Beyotime Biotechnology, China) lysis solution via a high-speed homogenizer machine. Incubate the homogenate on ice for 20 min, and next, all the homogenates were centrifuged at 13,400 rpm for 10 min at 4°C. Approximately 75% of the RIPA buffer volume was collected supernatant and thoroughly mixed after adding an equal volume of 2 Laemmli sample buffer. Each sample’s total protein concentration was detected using BCA assay kits (Pierce, Rockford, IL, USA). Equal amounts of protein were added to each lane and run on a 12 % SDS-PAGE gel at 18 V/cm to separate the total protein and transfer protein into PVDF membranes using a semi-dry transfer device (Bio-Rad, CA, USA). The membranes were collected and blocked for 1 hour at room temperature with 5% bovine serum albumin (BSA). Primary antibodies were used to incubate the membrane, with dilutions based on the ratios of the fellows: Sox9 (1:1000), Caveolin1 (1:500), and AR (1:500). In the refrigerator, each membrane was incubated with one primary antibody overnight in a shaker at 4°C in the refrigerator. The target protein was detected with the corresponding secondary antibodies after incubating the membrane for 1 h at a temperature. Finally, the data of target protein expression was normalized by endogenous β-actin and calculated via computing the ratio of the target protein of interest relative to the β-actin.

### Statistical analysis

For all statistical analysis of the data in this study, GraphPad Prism, Version 8.0 software was used. Data results were expressed as mean ± SD, and statistical methods were used for unpaired t-test. In the case of no special description, the significance is expressed as follows: n.s., not significant; *P< 0.05; **P < 0.01; ***P < 0.001; ****P < 0.0001, significant.

## Results

### Anatomic comparison of different developmental stages of the musk gland

By dissecting the musk gland tissue of Chinese forest musk deer (FMD) at different periods, the results show that the musk gland is already developing at 4 months of age (Supplementary Data S1 A), but the well-developed musk sachet is only present in adults (6-years-old) male FMD (Supplementary Data S1 B). Through the results of tissue anatomy, we compared the musk gland of developing (4 - month-old) and adult (6 years old) male FMD. In order to further study the correlation between skin and musk gland, and the developing FMD (4- month-old) was selected as the research object.

### Transcriptome sequencing data quality control

The transcriptome sequencing results of the musk gland, back skin, and abdominal skin are shown in Table 2. A total of 55, 237, 098 raw data of musk glands, 53, 488, 432 valid data, 96.83% data efficiency; skin-abdomen original data 54, 709, 886, valid data 49, 022, 732, data efficiency is 97.76%; the original data of back skin were 46, 598,196, and the valid data were 44, 357, 912, with a data efficiency rate of 95.19%. The percentages of Q20, Q30, and GC can reflect the quality of the clean reads. Quality data from musk gland and skin tissues reached over 97%, and the data from the other 30% also reached over 93%. This shows that the sequencing process was accurate and that the data came from high-quality sources.

**Table 2.**
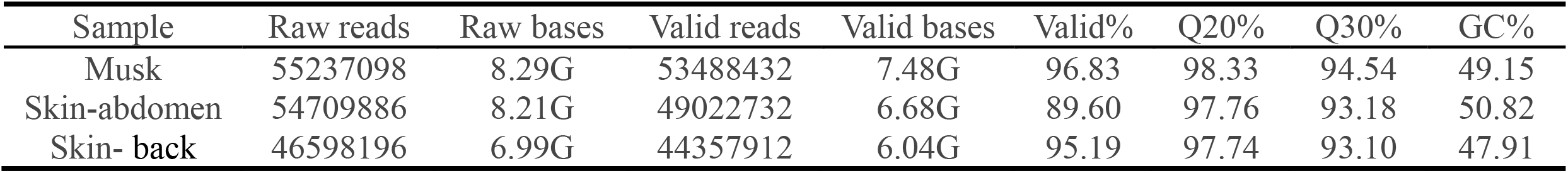
Overview of transcriptome sequencing data quality control

| Sample | Raw reads | Raw bases | Valid reads | Valid bases | Valid% | Q20% | Q30% | GC% |
| --- | --- | --- | --- | --- | --- | --- | --- | --- |
| Musk | 55237098 | 8.29G | 53488432 | 7.48G | 96.83 | 98.33 | 94.54 | 49.15 |
| Skin-abdomen | 54709886 | 8.21G | 49022732 | 6.68G | 89.60 | 97.76 | 93.18 | 50.82 |
| Skin- back | 46598196 | 6.99G | 44357912 | 6.04G | 95.19 | 97.74 | 93.10 | 47.91 |

The length distribution of assembled unigenes is shown in figure 1. The unigene length distribution reveals more genes in the range of 200-300bp, and more than 2000bp. The length distribution results for the analysis of these assembled gene sequences represent transcripts used for further analysis.

**Figure 1.**
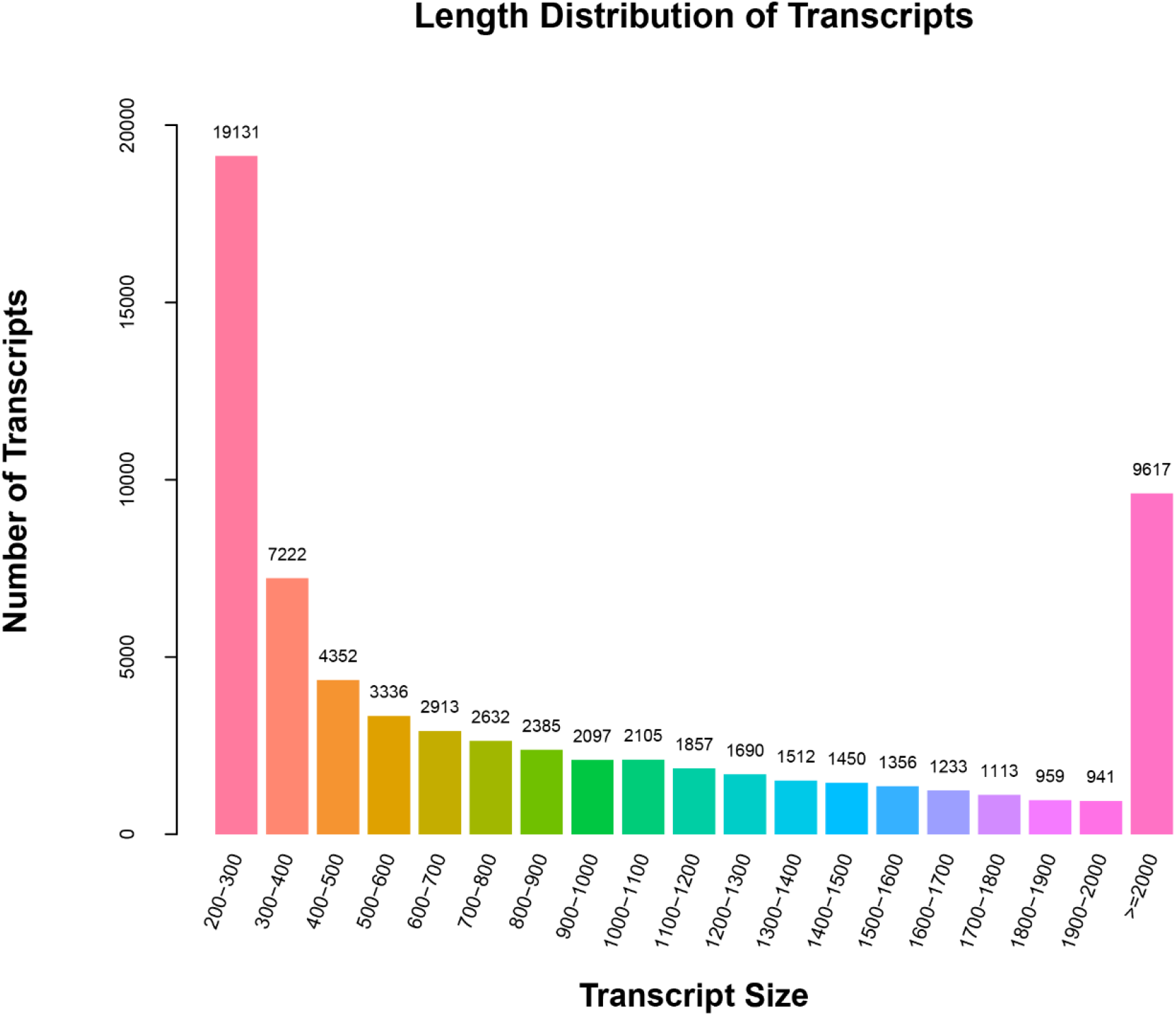
The unigenes length distribution of FMD. The x-axis indicates the length of sequenced unigenes, and the y-axis indicates the number of sequenced unigenes.

### Differential expression analysis of co-expressed genes

The number of genes co-expressed and specifically expressed between the three tissues of back skin, abdominal skin, and musk glands was analyzed using Venn diagrams, and the result is presented in figure 2. The vast majority of all genes (29, 771) in the back skin, abdominal skin, and musk gland tissues were expressed in all three tissues, accounting for 88.24% of the number of all predicted genes, with only a few genes having tissue specificity. Because most of the genes were co-expressed among the three tissues, musk glands, abdominal skin, and back skin tissues have a high degree of similarity at the genetic level. With the TPM value of genes in musk gland transcriptome greater than 25 as the screening condition, 2866 genes were obtained. Human genes were used as the background gene set, and gene enrichment analysis software metascape (https://metascape.org/gp/index.html#/main/step1) was used to analyze the cell and tissue specificity of the musk gland tissue (Zhou *et al*. 2019). The results of the Metascape analysis showed that these genes were significantly enriched in skin tissues (Figure 3). Excitingly, we analyzed abdominal skin, back skin, and using the same screening conditions described above, and obtained the same results that these genes were enriched in adipocytes, bronchial epithelial cells, and skin tissue (Supplementary Data S2 A-B). These results indicate a strong correlation between musk gland tissue and skin tissue.

**Figure 2.**
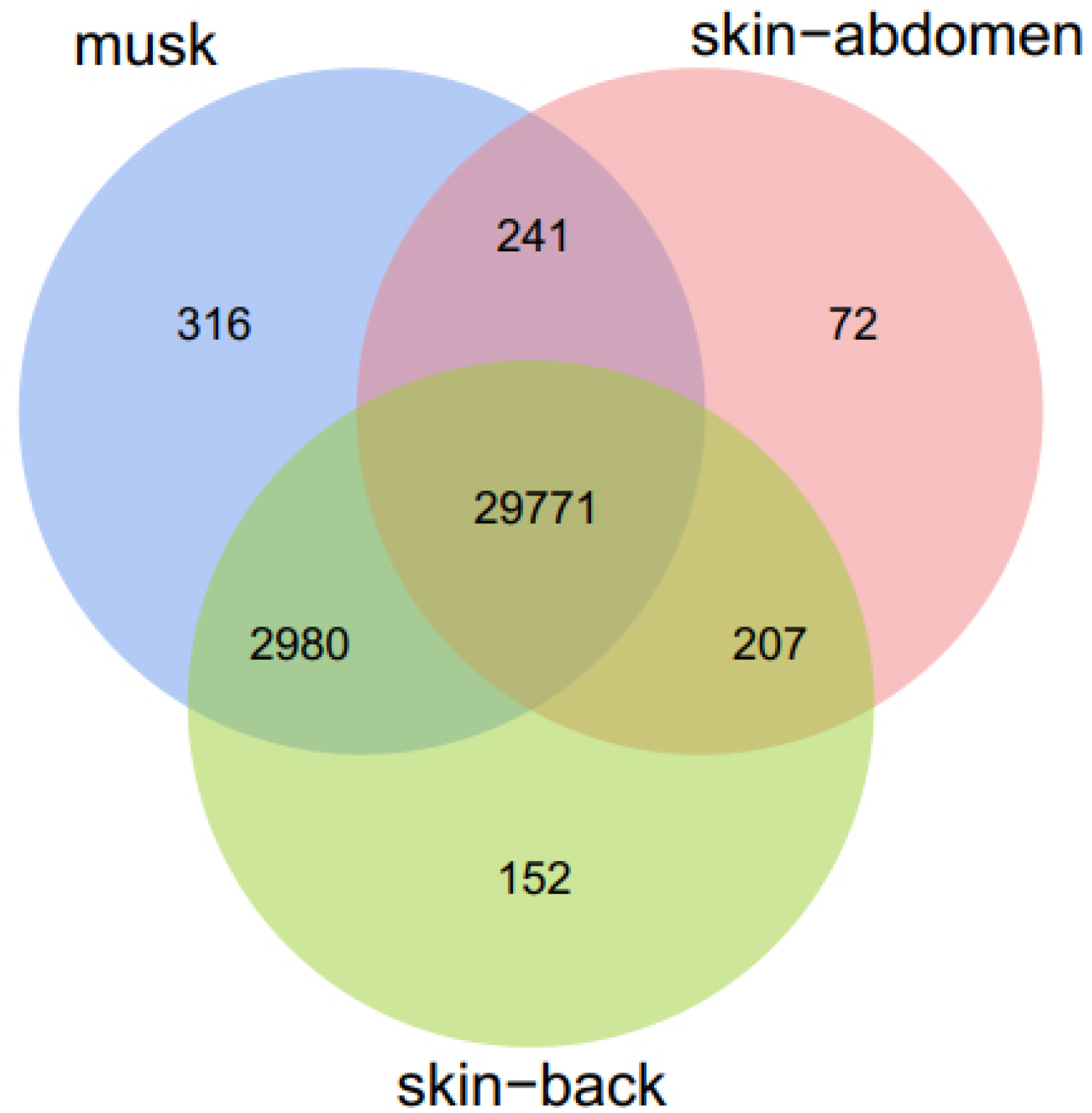
Venn diagram of genes in back skin, abdominal skin, and musk glands tissue. The Venn diagram illustrates the back skin, abdominal skin, and musk glands transcriptome datasets. The overlapping sections indicate the genes identified in both datasets.

**Figure 3.**
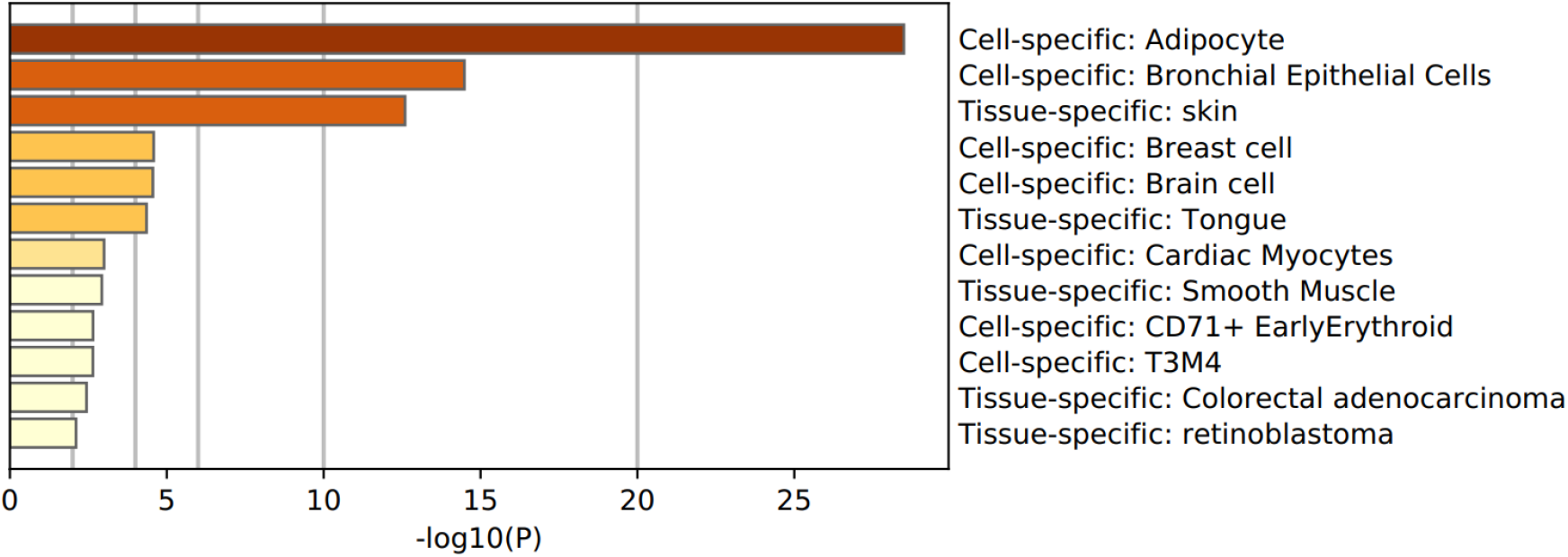
Gene metascape enrichment analyzed in FMD musk glands. The metascape software was used to analyze gene specificity in tissue and cells.

### Functional annotation of the transcriptome

The GO database is a database established by the Gene Ontology Consortium that contains a large amount of information on gene function. Using the GO database, genes and gene products can be annotated categorically into three categories: Biological Process, Molecular Function, and Cellular Component. Using a *P* value of less than 0.05 as a screening condition, the entries with the highest significance rank in each category in each group of GO enrichment results were presented based on the results of GO enrichment analysis of co-expressed genes in back skin, abdominal skin, and musk gland tissues (figure 4). 2, 803 genes were significantly enriched for 621 GO available entries in the analysis of the musk gland compared to the back skin, of which 155 were enriched to molecular functions, 393 for biological processes, and 73 for cellular composition. Significantly enriched pathways for structural constituents of muscle, focal adhesion, and adenylate cyclase-inhibiting dopamine receptor, respectively. In addition, entries related to skin development and differentiation, endodermal cell differentiation, and sebaceous gland cell differentiation (Supplementary Data S3 a). The analysis of the musk gland compared to the abdominal skin resulted in 4, 911 genes significantly enriched for 469 GO available entries, of which 79 were enriched for molecular functions, 316 for biological processes, and 74 for cellular composition. Enrichment to the most significant entries in these three categories: structural molecule activity, the establishment of protein localization, and extracellular exosome, respectively. In addition, there are functions related to endodermal cell differentiation and cytokine secretion (Supplementary Data S3 c). In the analysis of back skin compared to abdominal skin, 4, 435 genes were significantly enriched for 414 GO available entries, of which 90 were enriched to molecular functions, 263 for biological processes, and 61 for cellular composition (Supplementary Data S3 c).

**Figure 4.**
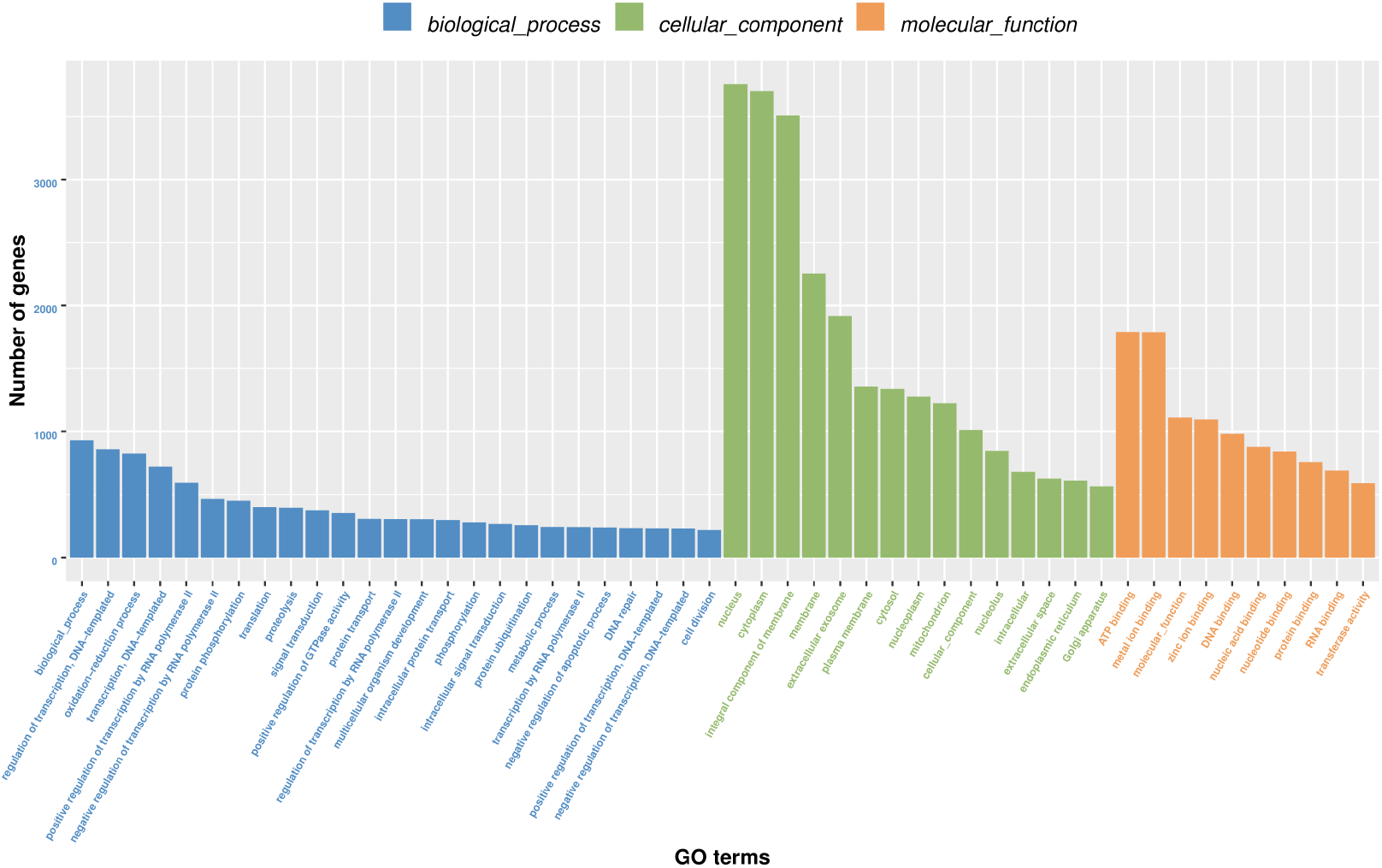
GO functional classification of co-expressed genes. The GO database allows for the annotation of genes and gene products in three categories: Biological Process, Molecular Function, and Cellular Component.

### KEGG signaling pathway prediction

We performed experimental KEGG pathway analysis to identify further the functional pathways of co-expressed genes in the abdominal skin, back skin, and musk glands. Prediction of transcriptomic data signaling pathways from abdominal skin, back skin, and musk gland tissues were divided into six main categories: biological systems, metabolism, human diseases, genetic information processing, environmental information processing, and cellular processes (Figure 5). A total of 1, 195 genes were significantly enriched for 99 KEGG metabolic pathways in the analysis of musk glands compared to back skin. The top three pathways with the highest significant enrichment levels were fat digestion and absorption, Glycerolipid metabolism, and Estrogen signaling pathway (Supplementary Data S4 a). A total of 1, 612 genes were significantly enriched for 55 KEGG metabolic pathways in the analysis of musk glands compared to abdominal skin, with the top three pathways with the highest significant levels of enrichment being Thermogenesis, Alzheimer’s disease, and Ribosome (Supplementary Data S4 b). A total of 1, 795 genes were significantly enriched for 68 KEGG metabolic pathways in the analysis of back skin compared to abdominal skin (Supplementary Data S4 c). The top three pathways with the highest significant enrichment levels were Thermogenesis, the Estrogen signaling pathway, and Huntington’s disease.

**Figure 5.**
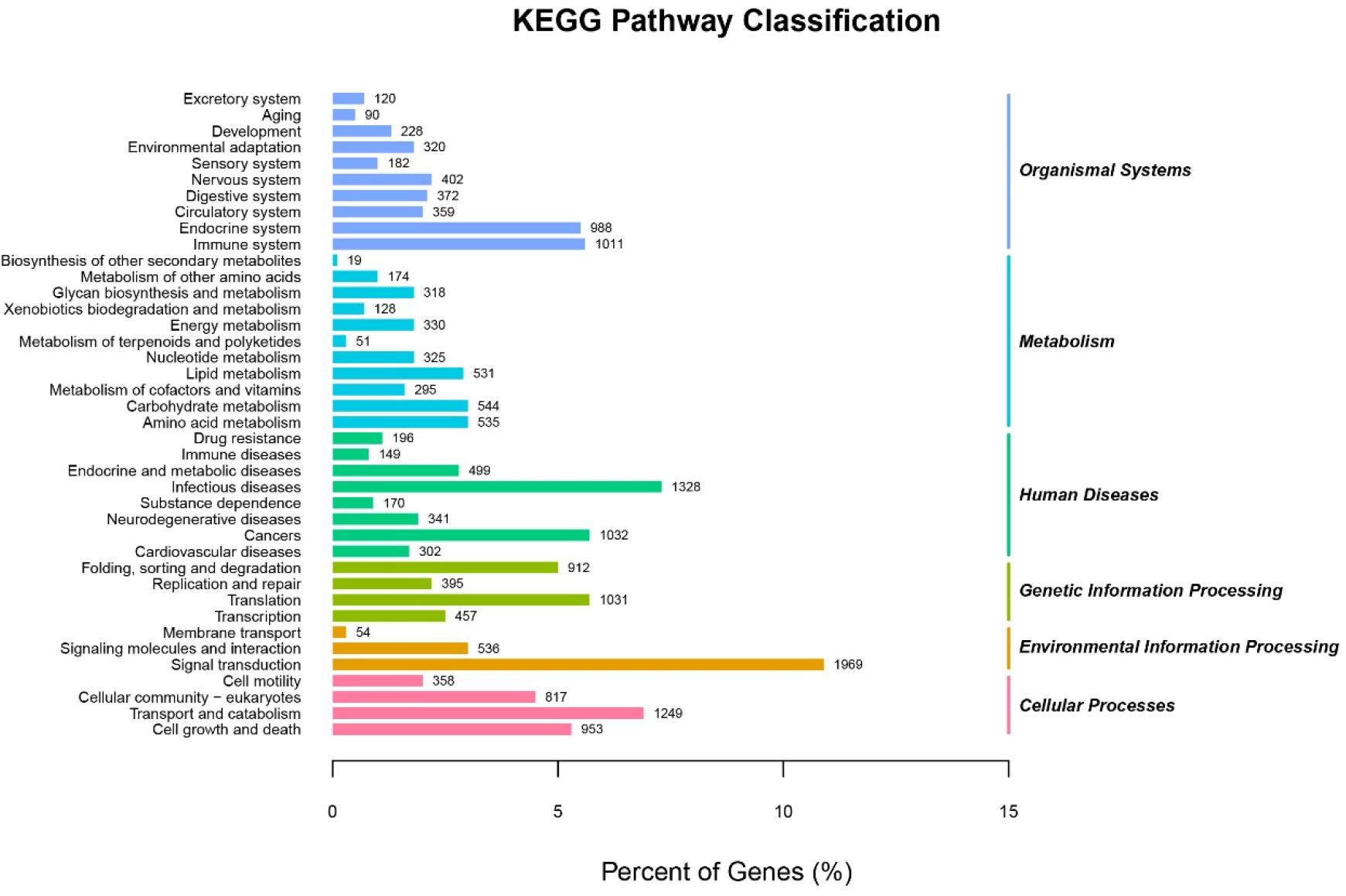
Histogram of KEGG pathway classification of co-expressed genes in abdominal skin, back skin, and musk glands tissue. The horizontal axis indicates the percentage of genes annotated to the pathway. The left vertical axis indicates the KEGG pathway, with different colors corresponding to the six categories in the top layer.

### The cellular structure of the different tissues in the musk glands

The cellular structure of the different tissues in the musk glands can be clearly visualized by HE tissues staining sections. The different tissues of the FMD musk gland, including the lax connective tissue (LCT) distributed therein sebaceous glands (SG), musk gland part (MGP), and acinar cavity (AC) (Figure 6A). The results of HE staining of the skin and hair follicles at the musk gland-skin junction showed that it had an intact hair follicle structure and abundant sebocytes (Figure 6B). The results of HE staining in the musk glandular region of FMD showed that the area possessed a large number of acinar cavity (AC) (Figure 6C). In order to further study the skin tissue, the longitudinal section and cross-section of the skin tissues used to perform HE staining. The longitudinal section structure of the skin tissue shows that from the inside to the outside are muscle tissue (MT), adipose tissue (AT), subcutaneous tissue (ST) and epidermis layer (EL) (Figure 6D). The results of cross-section of skin and hair follicle structure showed that the hair follicles surrounded by sebaceous glands at different stages of development are different. There are a large number of sebocytes in the early stage of development, but the hair follicle space is small, and then the well-developed hair follicles have few sebocytes around them, but there is a larger space around it (Figure 6E). The results of a complete hair follicle cross-section showed that it contains the hair shaft cuticle (Cut), the hair shaft cortex (Cor), and the hair shaft medulla (Med) (Figure 6F). In response to the phenomenon that hair follicles have different sebocytes encapsulations at different developmental stages, we conducted an in-depth study of the skin-hair follicle junction. The skin-follicles junction longitudinal section and HE staining was performed, and the results showed that sebaceous glands and hair follicles may act as a structural unit, including hair follicles (HF), sebaceous glands (SG), and hair shafts (HS) (Figure 6G), and the hair follicle is surrounded by a large number of sebaceous cells, marked by black arrows in the figure (Figure 6H). Figure 6I HE staining shows that a complete hair shaft structure, from inside to outside, the hair shaft medulla (Med), the hair shaft cortex (Cor), and the hair shaft cuticle (Cut) are respectively.

**Figure 6.**
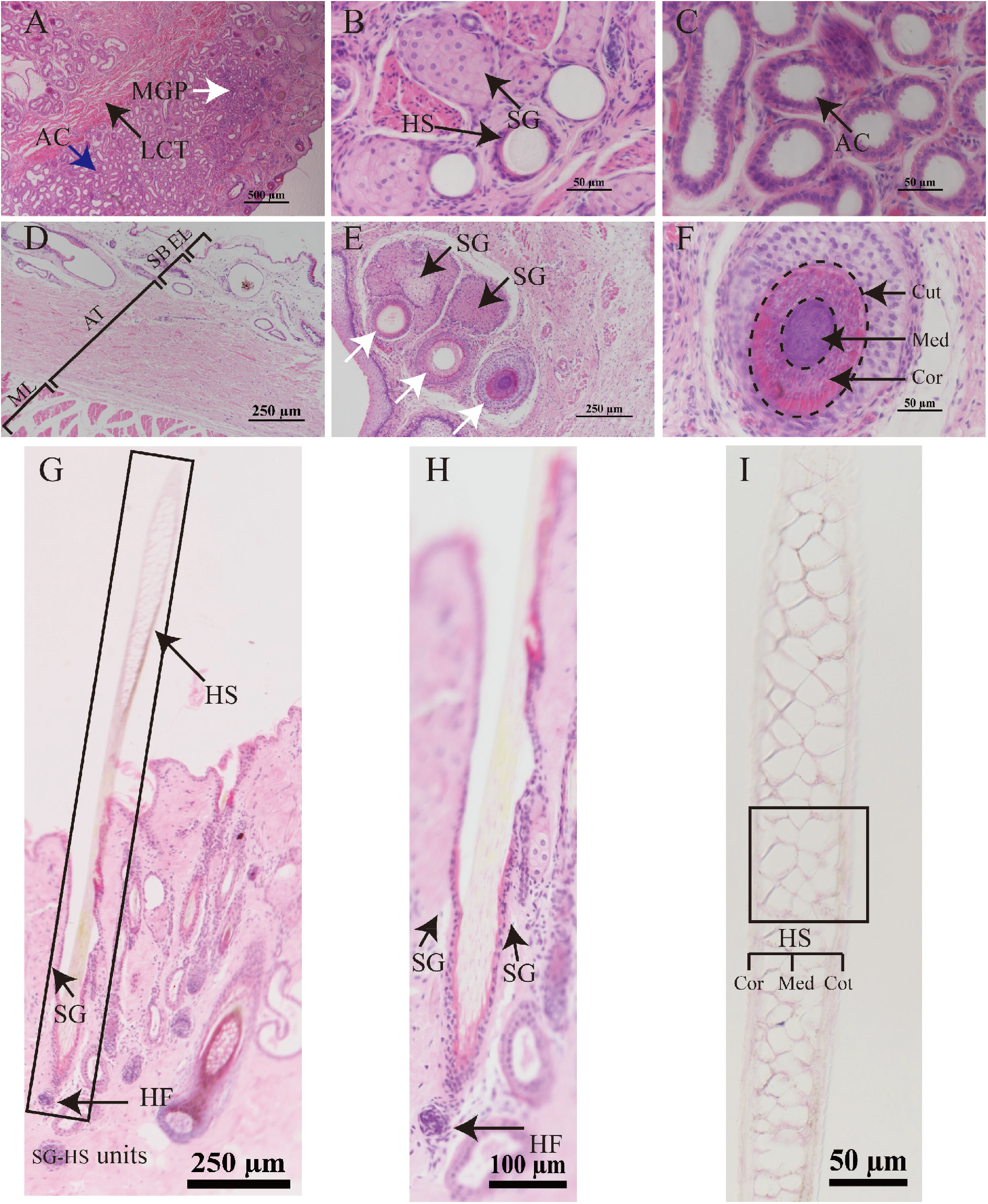
HE staining of musk glands and skin tissue of four-month-old FMD. (A) The general structure of the musk glands (4X objective lens; scale=500μm). The black arrow indicates the loose connective tissue (LCT), musk glands part (MGP), and the blue arrow indicates acinar cavity (AC); (B) The musk glands skin junction (40X objective lens; scale=50μm). The arrow points to the hair structure (HS) and sebaceous glands (SG); (C) The gland part (40X objective lens; scale=50μm). Blank space is the acinar cavity (AC); (D) The longitudinal section and HE staining of FMD skin tissue (10X): The figure shown the epidermis layer (EL), subcutaneous tissue (ST), adipose tissue (AT) and muscle tissue (MT) respectively. (E) The FMD skin tissue cross-section and HE staining. The black arrows indicate the sebaceous glands (SG) and white arrows refer to the three different developmental stages of hair follicle tissue; (F) The hair follicle structure of the FMD. The picture shows the hair shaft medulla (Med), hair shaft cortex (Cor), and hair shaft cuticle (Cut). (G) Forest musk deer skin tissue cross-section and HE staining. Rectangular box shows a structural unit of sebaceous gland and hair follicle (SG-HF unit), including sebaceous gland (SG), hair follicle (HF) and hair shaft (HS). (H). Hair follicle and sebaceous gland microstructure; (I) The complete hair shaft structure (HF), including hair shaft medulla (Med), hair shaft cortex (Cor), and hair shaft cuticle (Cut).

### Localization of Sox9, Caveolin1, and AR protein in musk gland

After immunohistochemical, the corresponding Sox9, Caveolin1, and AR protein localization and expression in back skin, abdominal skin, and musk gland slices were observed under light microscopy. In the negative control results, the nuclei were light blue in color, and the cytoplasm and background were colorless (Figure 7A1-A3). The results of immunohistochemistry of Sox9, Caveolin1, and AR in the back skin, abdominal skin, and musk gland tissues of FMD showed that Sox9 (Figure 7B1 -B3) (black arrow), Caveolin1 (Figure 7C1 -C3) (black arrow), and AR were localized in the back skin, abdominal skin, and musk gland tissue (black arrow), but AR showed lighter coloration in the back skin and abdominal skin (Figure 7D1-D3). The immunohistochemistry results showed that Sox9 and AR were mainly expressed in the nucleus of the musk gland and skin tissues, while Caveolin1 was localized on the membrane structure of cells in the musk gland and skin tissues (Figure 7). Also, based on visual observation, SOX9,Caveolin1, AR staining intensity gradients are described (table 3).

**Table 3.** the SOX9, Caveolin1, AR immunoreactivity based on visual inspection intensity

| Tissues | Protein |  |  |
| --- | --- | --- | --- |
|  | SOX9 | CAVEOLIN1 | AR |
| Skin-abdomen | ++ | ++ | ++ |
| Skin-back | +++ | +++ | +++ |
| Musk gland | +++ | ++ | +++ |

Immunoreactivity of tissue cell: moderate positive (++); strong positive (+++).

**Figure 7.**
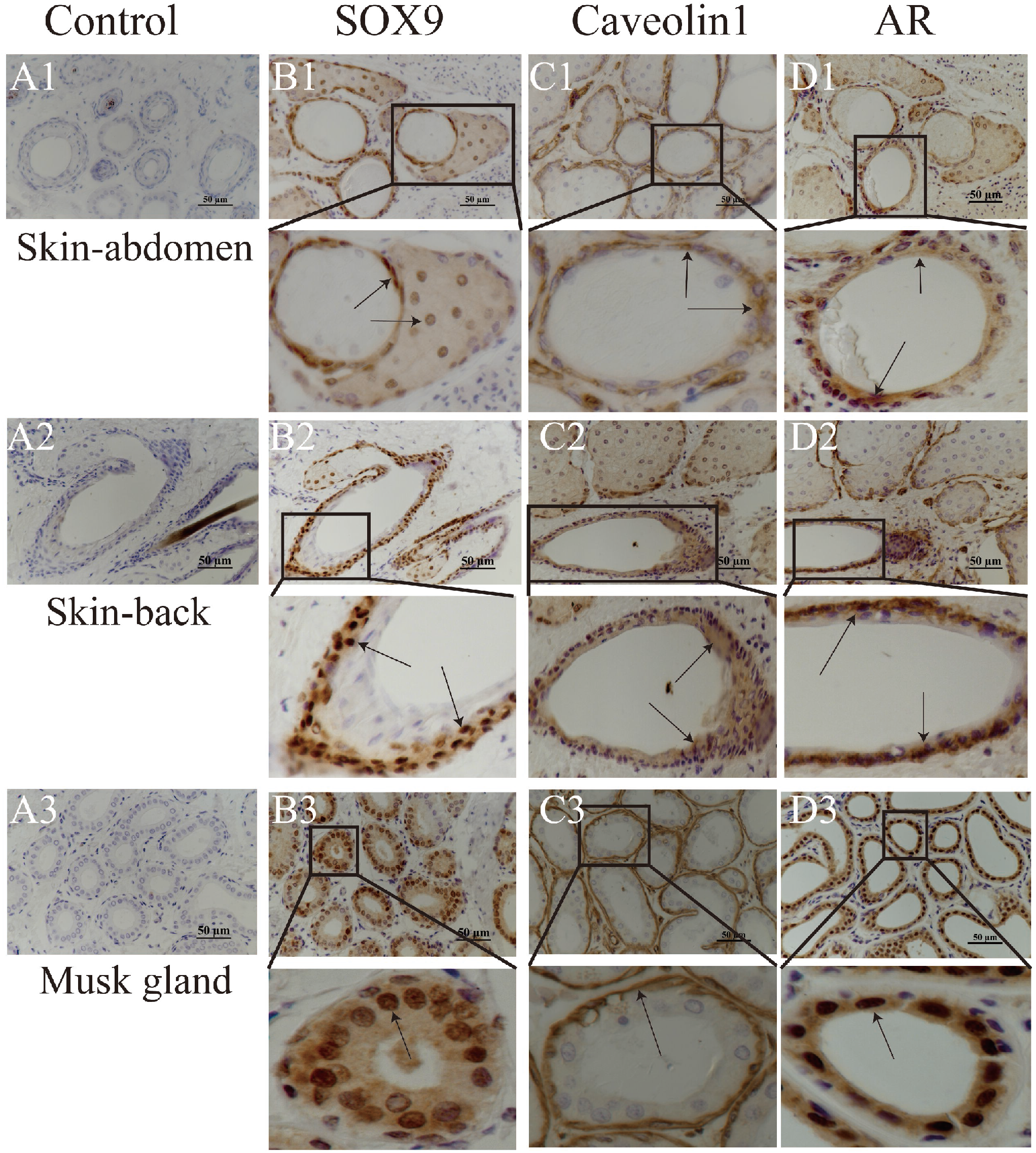
Negative control (A1-A2-A3). Protein expression and location of Sox9, Caveolin1, and AR proteins in abdominal skin (B1-C1-D1), back skin (B2-C2-D2), and musk glands (B3-C3-D3), respectively. Brown is the positive signal of the corresponding protein (40X objective lens; scale=50μm). Black arrows indicate positive signals.

### The *Sox9, Caveolin1* and *AR* mRNA levels were validated by RT-qPCR

To verify *Sox9, Caveolin1, AR* mRNA expression, we performed RT-qPCR to verify the expression profiles of the three genes in back skin, abdominal skin, and musk gland. The RT-qPCR results showed that *Sox9, Caveolin1, AR* were expressed in back skin, abdominal skin, and musk gland. In RT-qPCR results, Sox9 gene expression was highest in musk glands and lowest in abdominal skin. (Figure 8A). Similarly, *Caveolin1* expression was highest in the musk gland tissue and lowest in the abdominal skin (*p*< 0.05) (figure 8B). In addition, *AR* expression was the highest in the musk gland (*p*< 0.05), with non-significant differences in expression in the abdominal skin and back skin (*p*> 0.05) (figure 8C).

**Figure 8.**
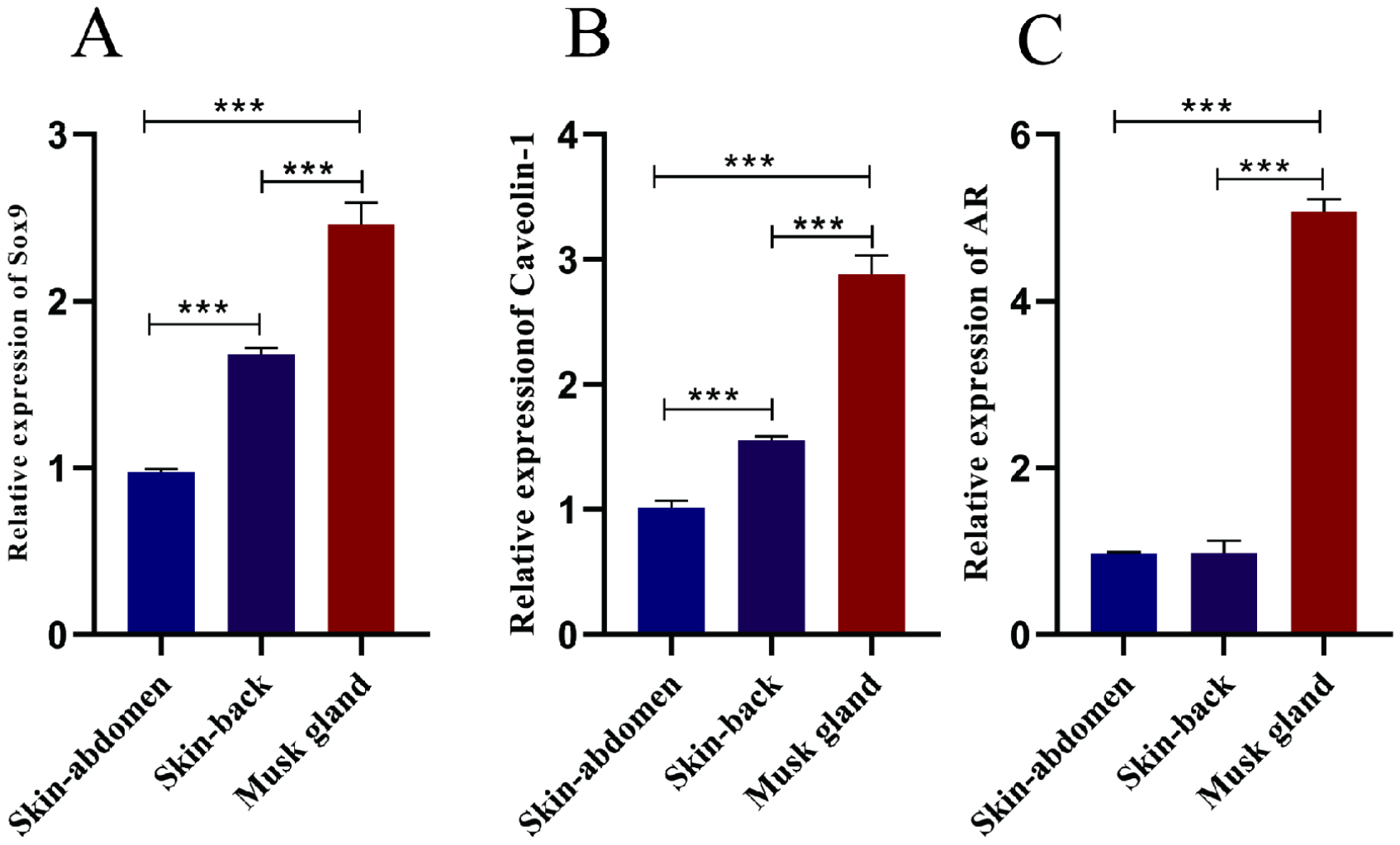
Expression of *Sox9, Caveolin1, AR* in abdominal skin, back skin, and musk glands was detected by RT-qPCR. The mRNA expression levels of *Sox9, Caveolin1, AR* were normalized by the mRNA expression levels of GAPDH. All data have three replicates, and the results are presented as mean ± SD. The bars chart superscripts indicate significant differences (*P *<* 0.05, **P *<* 0.01, ***P *<* 0.001, ****P < 0.0001), and an unpaired t-test was used for analysis.

### Expression of Sox9, Caveolin1, and AR protein

To further evaluate the role of Sox9, Caveolin1, and AR in the development of abdominal skin, back skin, and musk glands, protein expression was detected using western blotting. The results are presented in Figure 9. Figures 9A and 9B showed that Sox9 was most highly expressed in the musk gland and least expressed in the abdominal skin. Caveolin1 and AR are highly expressed in the musk gland (*p*< 0.05) and are not statistically different in the abdominal skin and back skin (*p*> 0.05) (Figures 9C-D).

**Figure 9.**
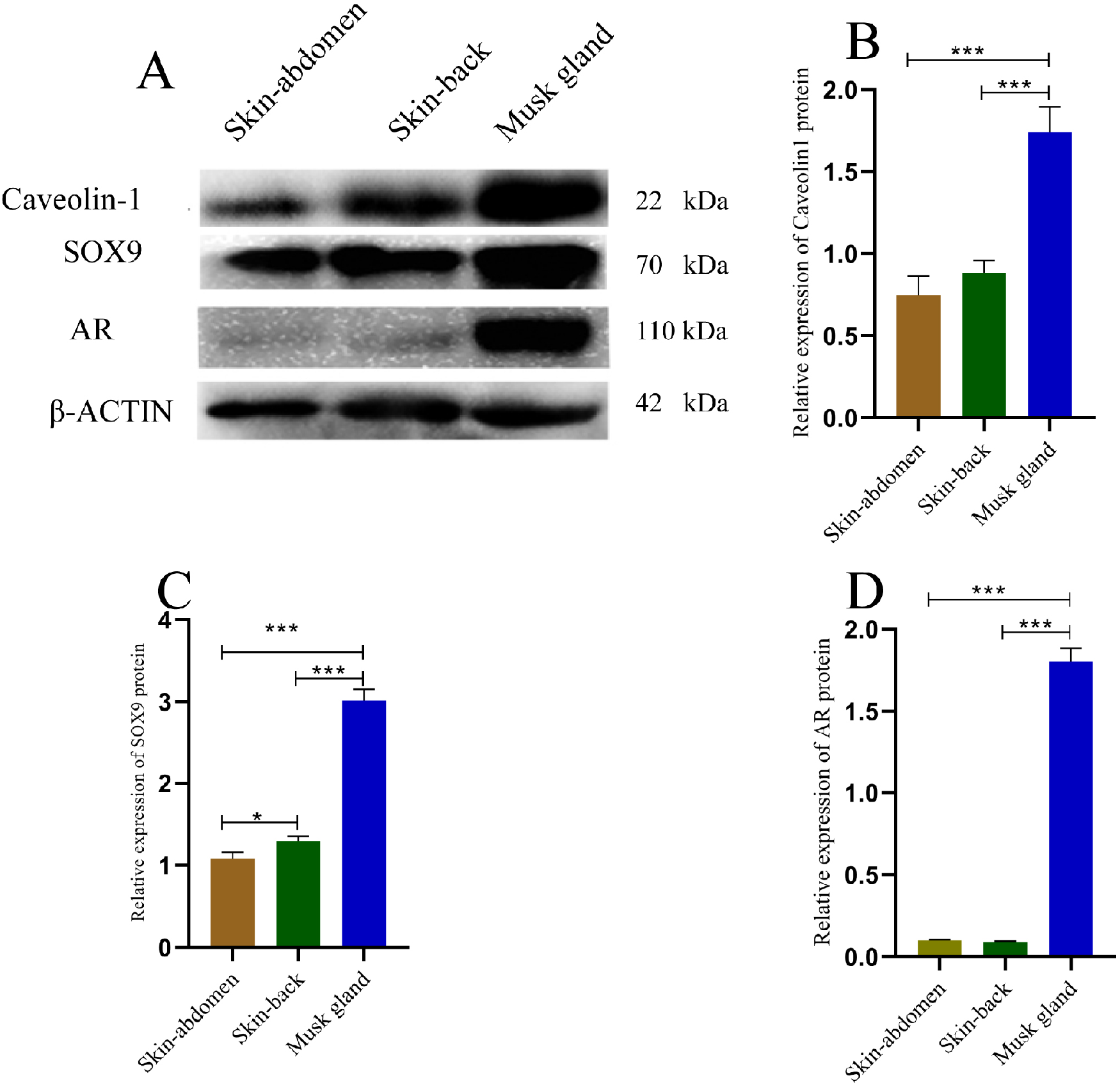
Protein expression of Sox9, Caveolin1, and AR in abdominal skin, back skin, and musk glands tissue. A. Western blotting for detecting Sox9, Caveolin1, AR protein expression, and β-actin as an internal reference for protein detection. B-D. All samples have three replicates, and the results are presented as mean ± SD. The bars chart superscripts indicate significant differences (*P *<* 0.05, **P *<* 0.01, ***P *<* 0.001, ****P < 0.0001), and an unpaired t-test was used for analysis.

## Discussion

Musk is a special secretion secreted by adult males with a strong and distinctive scent signatures that can be used to mark their territory and attract females (Zhang *et al*. 2021a). Fatty acid oxidation and related metabolic pathways are necessary for musk secretion in FMD by generating large amounts of energy (Zhang *et al*. 2021b). The sebaceous glands of the FMD musk gland are involved in musk secretion (Thody and Shuster 1989; Downie *et al*. 2004; Smith and Thiboutot 2008; Zouboulis *et al*. 2008; Schneider and Paus 2010; Chen *et al*. 2018). *Sox9, Caveolin1*, and *AR* all play an important role in regulating the lipid metabolism of the sebaceous glands (Bastiani *et al*. 2009; Barrault *et al*. 2015; Davey and Grossmann 2016; Shi *et al*. 2017). However, there is a blank in studies on relevance of *Sox9, Caveolin1*, and *AR* in the developmental process of FMD skin tissues and musk gland tissue. Both skin tissues and musk gland tissue have sebum secretion functions. Therefore, we hypothesized that there is some similarity between skin tissue and musk gland tissue. *Sox9, Caveolin1*, and *AR* play an important role in regulating sebum secretion and cell proliferation. Therefore, we will explore the correlation in the skin tissues and musk gland of the developing male FMD by RNA-seq, immunohistochemistry (IHC), western blotting (WB), and RT-qPCR.

Through the dissection of different periods of male forest musk deer (FDM), it is clear that the well-developed musk gland sachet only exists in adult male FMD (Supplementary Data S1 B). The musk gland is a unique organ of FMD, which can synthesize and secrete musk, and it is synchronized with the development of testis (Fuchs 2016). As previously reported, adult male FMD can secrete musk during the breeding season, and the secretions need to enter the musk gland sachet, and immature musk in the capsule takes about two months to mature (Zhang *et al*. 2017; Chen *et al*. 2018; Lv *et al*. 2022). The four-month-old forest musk gland is still developing, and at the same time, it does not have the function of secreting musk, nor does it lack the well-developed musk gland sachet that contains musk (Supplementary Data S1 A). However, only adult males can secrete musk and have the well-developed musk gland sachet to storage and maturation (Supplementary Data S1 B).

For a further and more comprehensive understanding of the correlation between skin tissues and musk gland, we analyzed their transcriptome data. The Q20% of musk gland and skin tissue reached more than 97% and the Q30% reached more than 93% (Table 2). It indicates the high precision of detection and high quality of sequencing data during sequencing (Andrews 2014). The unigenes assembly results showed a high number of genes of 200-300 bp and more than 2000 bp (Figure 1).

Based on previous studies, it has been shown that these assembly results can be used for further analysis (Zhou *et al*. 2019). The genes co-expressed and specifically expressed in the three tissues of the abdominal skin, back skin and musk gland were analyzed by Venn diagram. Venn diagram analysis showed that genes co-expressed in abdominal skin, back skin, and musk gland tissues accounted for 88.24% (29,771) of the total genes (Figure 2). The Metscape database, as an open system, provides a systematic data analysis platform for bioinformatics analysis (Zhou *et al*. 2019). The results of the analysis using Metascape showed a strong correlation between the musk gland and the skin (Figure 3). Genes and gene products can be classified and annotated into three categories: biological processes, molecular functions, and cellular components, which are analyzed through the GO database (Figure 4). KEGG analysis of the pathways involved in the musk glands’ metabolic and synthetic activities and skin tissues was annotated, including biological systems, metabolism, human diseases, genetic information processing, environmental information processing, and cellular processes (Figure 5). Regarding the transcriptome data, in general, up to 88.24% of the genes co-expressed in the abdominal skin, back skin, and musk gland tissues, it is speculated that the three have a strong correlation. The sebaceous glands of the skin tissue are spatially correlated with the hair follicles and the musk glands in terms of histological structure (Figure 6B-E). Interestingly, the sebaceous gland lines wrapped around hair follicles at different development stages are different, and the space around hair follicles is also different (Figure 6E). Early hair follicles are surrounded by a large number of sebocytes, but the space around the hair follicle is small, while the well-developed hair follicle has few sebocytes around the hair follicle, but the space is larger. This may suggest that only well-developed hair follicles can excrete the secretions of glands such as sebaceous glands. This result is consistent with previous studies on the correlation between hair follicles and skin development (Schneider *et al*. 2009). In response to this exciting finding, we conducted experiments on the skin and hair follicle structure from a different perspective, that is, longitudinal section of skin and hair follicle junction. Sebaceous glands and hair follicles are closely connected in the spatial structure (Figure 6G-I), and may participate in the secretion of sebum as a functional structural unit. The sebaceous glands may form sebum units with hair follicles and reach the skin surface via the hair canal, and play various functions such as antibacterial, antioxidant and pheromone communication (Ehrmann *et al*. 2016; Schneider and Marlon 2016; Schneider and Zouboulis 2018).

In order to further understand the development of the abdomen skin, back skin, and the musk gland tissue. Based on the transcriptome data and previous related studies, we chose *Sox9, Caveolin1*, and *AR* as the research objects, through immunohistochemistry (IHC), western blotting (WB), and RT-qPCR. Previous studies have shown that *Sox9* has multiple functions in cell fate regulation, stem cell behavior, embryonic development, acquired and congenital diseases (Jo *et al*. 2014). *Caveolin1* plays a key role in regulating the proliferative capacity of epidermal stem cells, altering the proliferative capacity of epidermal stem cells and promoting the healing of wounded skin (Yang *et al*. 2019). Androgens are primarily synthesized in the testes and have long been thought to be male sex steroids involved in male characteristics and androgens mainly performed their functions via *AR* (Rege *et al*. 2013; Storbeck *et al*. 2013; Pretorius *et al*. 2016). In addition, the *AR* is seasonally expressed in the epididymis of the FMD (Liu *et al*. 2019), which means that it plays an important role during the breeding and non-breeding seasons. IHC results showed that Sox9 and AR were localized in the nucleus and Caveolin1 in the cytoplasm in the abdominal skin, back skin, and musk gland (figure 7). The RT-qPCR results showed that *Sox9, Caveolin1*, and *AR* were highly expressed in the musk gland tissue (figure 8). Similarly, WB results also showed that Sox9, Caveolin1, and AR were highly expressed in the musk gland tissue (figure 9). Sox9 positive cells are considered to be a population of multipotent stem cells with the ability to differentiate into sebaceous gland cells and epithelial cells (Horsley *et al*. 2006; Nowak *et al*. 2008; Qiu *et al*. 2014). Caveolin1 is involved in regulating cell proliferation (Tamai *et al*. 2001; Faggi *et al*. 2015). The number of cells expressing stem cell maker was increased in several organs, including the intestine, mammary gland, and brain of Caveolin1 null mice (Li *et al*. 2005; Sotgia *et al*. 2005; Jasmin *et al*. 2009). *Caveolin1* may play a negative regulatory role in the proliferation of stem cells, such as neural progenitor cells bone marrow-derived mesenchymal stem cells (MSCs) (Park *et al*. 2000; Case *et al*. 2010; Lee *et al*. 2010; Samarasinghe *et al*. 2011; Baker *et al*. 2012). However, *Caveolin1* is able to promote cell proliferation in epidermal cells (Yang *et al*. 2019). This may be that Caveolin1 can participate in various protein interactions and is the key factor of cell signal transduction (Boscher and Nabi 2012). In the classical testosterone signaling pathway, androgen exerts its genomic effect through AR, and the testis mainly produces testosterone, and the development of the gland is synchronized with the testis (Quigley *et al*. 1995). Previous results showed that both *Sox9* and *Caveolin1* could interact with *AR*, while androgens play an important regulatory role in the secretion process of the musk gland (Bennett *et al*. 2009; Khurana and Sikka 2019). Together, these results suggest that there is a strong correlation between skin tissue and musk gland and *Sox9, Caveolin1*, and *AR* may play important roles in skin and musk gland development and sebum secretion.

In summary, this study compared and analyzed the tissue morphology, gene expression, and protein expression of abdominal skin, back skin, and musk gland through HE, IHC, WB, RT-qPCR, RNA-seq. The co-expressed genes of abdominal skin, back skin and musk glands are as high as 88.24%, and the highly expressed genes are mainly enriched in adipocytes, bronchial epithelial cells and skin tissue, which implies an extremely strong correlation among the three tissues (Figure 10). The key regulatory genes, *Sox9, AR*, and the signal transduction hub gene *Caveolin1*, were expressed in the abdominal skin, back skin, and musk glands. Together, these data suggest that the musk glands and skin tissues have a strong correlation.

**Figure 10.**
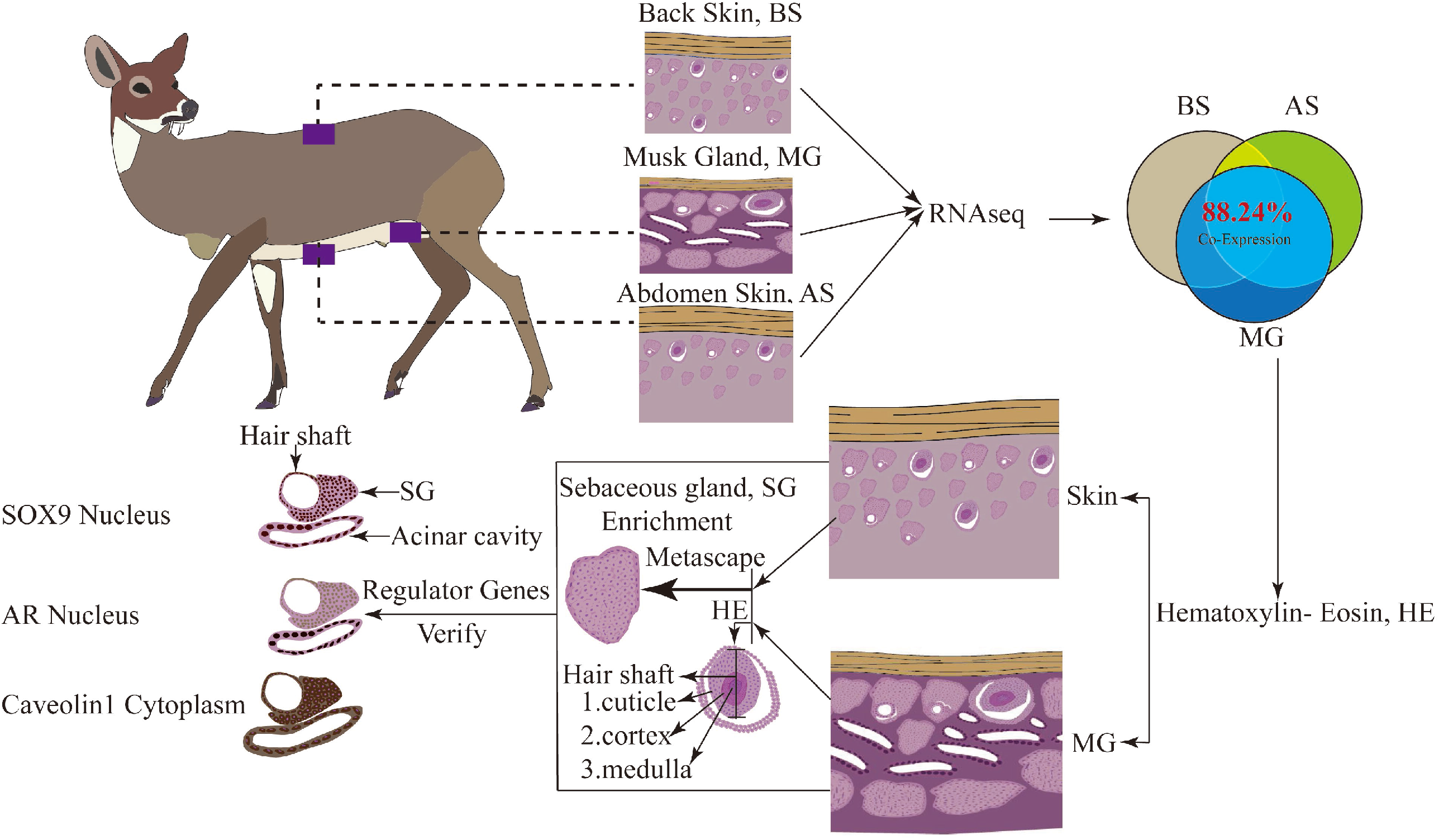
The schematic representation of the correlation in abdominal skin, back skin and musk glands. This figure explains the correlation of back skin, abdominal skin and musk glands in tissues, cells and genes of FMD.

## Data availability

All data in the study is available upon request. Supplementary Data S1 anatomical diagram of the development of the musk glands at different periods. Supplementary Data S2 gene metascape enrichment analyzed in FMD abdomen skin and back skin. Supplementary Data S3 GO enrichment analysis of differentially expressed genes in back skin, abdominal skin, and musk glands tissue. Figure a shown that the musk gland’s analysis result compared to the back skin. Supplementary Data S4. KEGG pathway enrichment analysis bubble chart.

Supplementary materials, which will be uploaded to the genetics website.

## Acknowledgments

All authors acknowledge Mr. Xuezhe Zhang at the Shaanxi Baosen Musk Deer Industry Co., Ltd. (Shaanxi, China) for providing experimental animals.

## Funding

This work supported by the Science and Technology Innovation Program of Shaanxi Academy of Forestry (SXLK2021-0219).

## Conflicts of interest

The author states that there is no conflict of interest.

---

**Supplementary Data S1**. Anatomical diagram of the development of the musk glands at different periods. Figure A shows the anatomy of the musk glands of a 4-mouth-old FMD; figure B shows the development of an adult (six-year-old) FMD musk sachet.

**Supplementary Data S2**. Gene metascape enrichment analyzed in FMD abdomen skin and back skin. Figure A shows the results of the abdominal-skin gene enrichment analysis. Figure B shows the results of the back-skin gene enrichment analysis.

**Supplementary Data S3**. GO enrichment analysis of differentially expressed genes in back skin, abdominal skin, and musk glands tissue. Figure a shown that the musk gland’s analysis result compared to the back skin. Figure b shown the musk gland’s analysis result compared to the abdominal skin. Figure c shown that analysis result of back skin compared to the abdominal skin.

**Supplementary Data S4**. KEGG pathway enrichment analysis bubble chart. Figure a shown the KEGG analysis result of the musk glands compared to the back skin. Figure b shown the KEGG analysis result of the musk glands compared to the abdominal skin. Figure c shown the KEGG analysis result of back skin compared to the abdominal skin.

